# Longitudinal Visual Analytics for Unpacking the Cancer Journey

**DOI:** 10.1101/444356

**Authors:** Zhou Yuan, Sean Finan, Jeremy Warner, Guergana Savova, Harry Hochheiser

**Affiliations:** University of Pittsburgh, Pittsburgh, PA, USA; Boston Children’s Hospital, Boston, MA, USA; Vanderbilt University, Nashville, TN, USA

## Abstract

Retrospective cancer research requires identification of patients matching both categorical and temporal inclusion criteria, often based on factors exclusively available in clinical notes. Although natural language processing approaches for inferring higher-level concepts have shown promise for bringing structure to clinical texts, interpreting results is often challenging, involving the need to move between abstracted representations and constituent text elements. We discuss qualitative inquiry into user tasks and goals, data elements and models resulting in an innovative natural language processing pipeline and a visual analytics tool designed to facilitate interpretation of patient summaries and identification of cohorts for retrospective research.

## Introduction

The complexities of cancer care create significant challenges for the extraction of information for retrospective research. As patients progress through diagnosis to treatment and subsequent monitoring, multiple encounters with varying specialists generate a rich set of clinical notes. For patients undergoing lengthy or multimodal (e.g., a combination of surgery, chemotherapy, and radiotherapy) treatment, hundreds or thousands of notes can be generated along the cancer journey. Review of these notes can be a laborious interpretive challenge, often involving many hours of time for medical professionals who must read through collections of notes to prepare summarized abstractions in spreadsheets or databases. This process is also brittle, as reviews conducted for one study may miss items of potential interest to subsequent studies. Although ad hoc solutions such as the oncologic history have spontaneously developed as information collection devices, they are not necessarily universal, accurate, or complete^1^.

The Cancer Deep Phenotype Extraction (DeepPhe) project is developing informatics solutions to overcome these inefficiencies. Unlike prior work applying Natural Language Processing (NLP) techniques to individual cancer documents^2–5^, DeepPhe combines details from multiple documents to form longitudinal summaries. Classic and state-of-the-art NLP techniques for extracting individual concepts are used alongside cross-document co-resolution techniques and inference rules and a rich information model^6^, to summarize diagnoses, treatments, responses and temporal relationships as needed to support retrospective research^7^.

Support for manual review of these results is critical. Visual and interactive tools help researchers interpret the complexities of relationships between cancers, tumors, treatments, responses, biomarkers, and other key attributes. Tools must support interpretation at multiple levels, moving from higher-level summaries to specific textual items informing those summaries. Tackling these challenges requires linking summary views with original text sources.

We draw upon a substantial body of prior work on visual cohort extraction tools, many of which have used temporal or flow metaphors to characterize temporal trends or transitions across patient populations^8–11^. DeepPhe visualization tools will extend these efforts with facilities for addressing challenges associated with the ambiguities of interpreting natural language.

We have developed a preliminary tool for visualization of patient summaries and identification of cohorts grouped by high-level concepts, e.g., cancer stage. The design of this tool was motivated by insights from qualitative inquiries with potential users and informed by a multi-level information model designed to bridge the gap between individual text spans and concepts relevant to domain experts.

## Methods

### Qualitative Inquiry

Interviews with cancer clinical researchers informed design of DeepPhe artifacts including an information model, an NLP pipeline, and visual analytics tools. Discussions focused on challenges in cancer retrospective research, including goals, information needs, representations, bottlenecks, and challenges. Information artifacts currently in use for managing abstracted retrospective were discussed. All interviews were audio-recorded. Interview notes and recordings were analyzed to extract information needs, problems, design suggestions, and other relevant information. Results from these analyses were used to develop user personae describing potential DeepPhe users, user stories describing specific tasks, and competency questions detailing specific information requirements. Subsequent to preliminary interviews, selected informants participated in card-sorting exercises focusing on identifying information items needed by researchers in specific domains. The University of Pittsburgh Human Research Protection Office classified the qualitative inquiries as exempt (PRO13120154).

### Information Model and Natural Language Processing Tool Development

Qualitative inquiry results were used to develop an information model capable of representing relevant items and attributes at multiple granularities, ranging from individual text mentions to patient summaries^6^. Existing cTAKES^12^ pipelines were extended to extract cancer information, to use rules to infer higher-level summaries, and to store results in a Neo4j graph database (www.neo4j.com). The initial DeepPhe architecture is described in detail by Savova, et al.^7^.

### Visualization

Insights from qualitative inquiries informed the development and iterative revision of a set of requirements and a corresponding series of low-fidelity prototypes for the visual analytics tools. As implementation of the prototype visualization proceeded in parallel with development of the NLP pipeline, speculative additions to the data model were constructed as needed to support visualization development for data items and attributes that were not fully supported by the NLP tool.

The resulting visualization tool was developed as a web application, using the Node.JS (www.nodejs.org) web platform to provide a middle-ware layer capable of retrieving data through the Neo4j REST interface. The visualization interface was implemented in HTML, CSS, Javascript, and the D3 visualization toolkit^13^.

## Results

### Qualitative Inquiry and Visual Analytics Requirements

User challenges identified during interviews involved difficulties with information availability, access, quality, and interpretation. Although some issues were specific to the types of cancer or the context of care, most were more broadly applicable (Table 1).

**Table 1:**
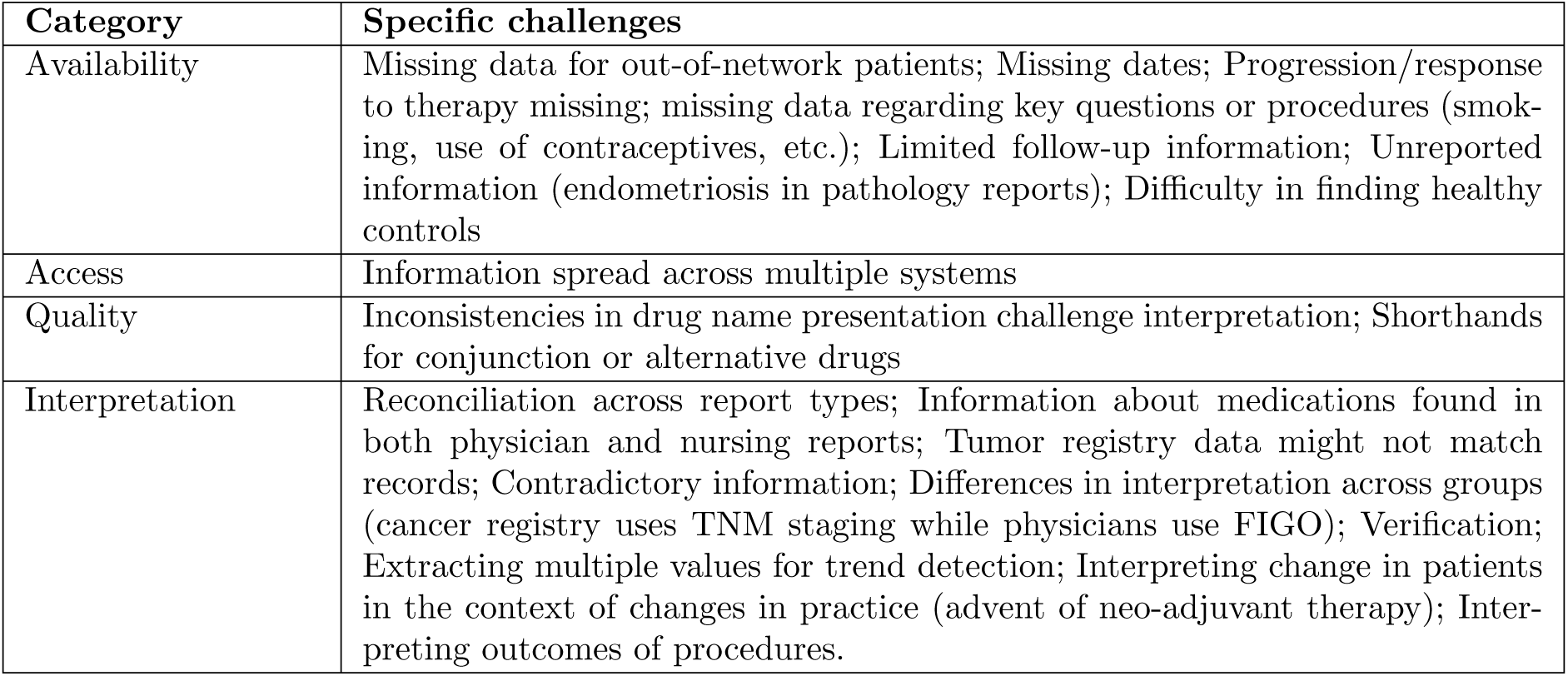
User information challenges identified during contextual interviews.

Together with informant descriptions of information needs and goals, these challenges informed the creation of more than 20 user stories detailing specific tasks to be conducted for individual patients and/or at the cohort level. These user stories were broadly grouped into 12 requirement categories (Table 2).

**Table 2:** System requirements and associated user stories

| Requirement | Granularity |  | User Stories |
| --- | --- | --- | --- |
|  | Cohort | Patient |  |
| R1: Overview | ✓ | ✓ | Explore distribution of available data |
| R2: Temporal | ✓ |  | Note time points or intervals corresponding to changes in clinical practice or other contextual factors that might influence results; Examine trends in lab values; Review sequences of care episodes; Ask temporal queries; Determine length of overall and progression-free survival |
| R3: Text |  | ✓ | Refer to original clinical text |
| R4: Provenance |  | ✓ | Identify original text used to extract/infer observations |
| R5: Multirecords |  | ✓ | Use multiple types of records to bolster interpretations |
| R6: QA |  | ✓ | Verify accuracy and relevance of extracted data; Check for consistency within documents set, and between documents and unstructured or registry data |
| R7: Stage |  | ✓ | Identify cancer stage |
| R8: Genomics |  | ✓ | Determine biomarker status and method used to make determination |
| R9: Treatments |  | ✓ | Identify treatments provided, number of cycles, doses, treatments offered but not taken, reasons for discontinuing treatments; Relate start and end dates of treatments to outcomes; Understand response to treatment/progression, including failure to respond, recurrence/no benefit, metastasis, developed resistance |
| R10: Filters | ✓ | ✓ | Filter cohorts based on categorical or temporal values; Focus patient exploration based on relevant constraints |
| R11: Uncertainty | ✓ | ✓ | Communicate both inherent ambiguity in notes and confidence in extracted information |
| R12: Search | ✓ |  | Conduct searches based on clinical criteria and temporal relationships |
| R13: Comparison | ✓ |  | Compare outcomes across selected cohorts |
| R14: Criteria | ✓ |  | Report criteria used to identify or compare cohorts |

### Information Model and Natural Language Processing Tools

Qualitative interview responses led to the development of an four-level information model for detailed cancer phenotypes, described previously^6^ and summarized here:

#### Mentions

Text spans in source documents discussing concepts and relations of interest, including tumors, body locations, treatments, stage indicators, biomarkers, and other key elements.

#### Compositions

Aggregation of all details from a given clinical note.

#### Episodes

Collection of documents into key event intervals, initially including work-up, diagnosis, medical decision making, treatment, and follow-up.

#### Phenotypes

Descriptions of cancers, tumors, treatments, and genomics, abstracted across the entire span of the patient history.

NLP tools for extraction of basic elements were implemented within the cTAKES environment^12^, with appropriate extensions for rules processing to support inference of higher-level summaries from text mentions. Distinct episode models were trained for breast, ovarian, and melanoma corpora. Additional details on the system architecture have been published previously^7^.

As development of the DeepPhe NLP tools is an ongoing effort, prototype implementation of the visual analytics tools has been facilitated by the construction of synthetic details to complete fields that cannot yet be extracted by DeepPhe. The current prototype displays extracted results for cancer stage, diagnosis, treatments, tumor size, histologic type, tumor extent, cancer cell line, body site, and biomarkers. Synthesized results for date of birth and menopausal status are also displayed.

### Visual Analytics Environment

Initial development of the information exploration tool has focused on a subset of requirements (R1:Overview, R2:Temporal, R3: Text, R7: Stage, and R8:Genomic) at the patient level:

*Patient details* (Figure 1A) are shown in a series of panels, beginning with patient demographics and summary cancer attributes, treatments, cancer stage, cell line, and TNM values^14^.

**Figure 1:**
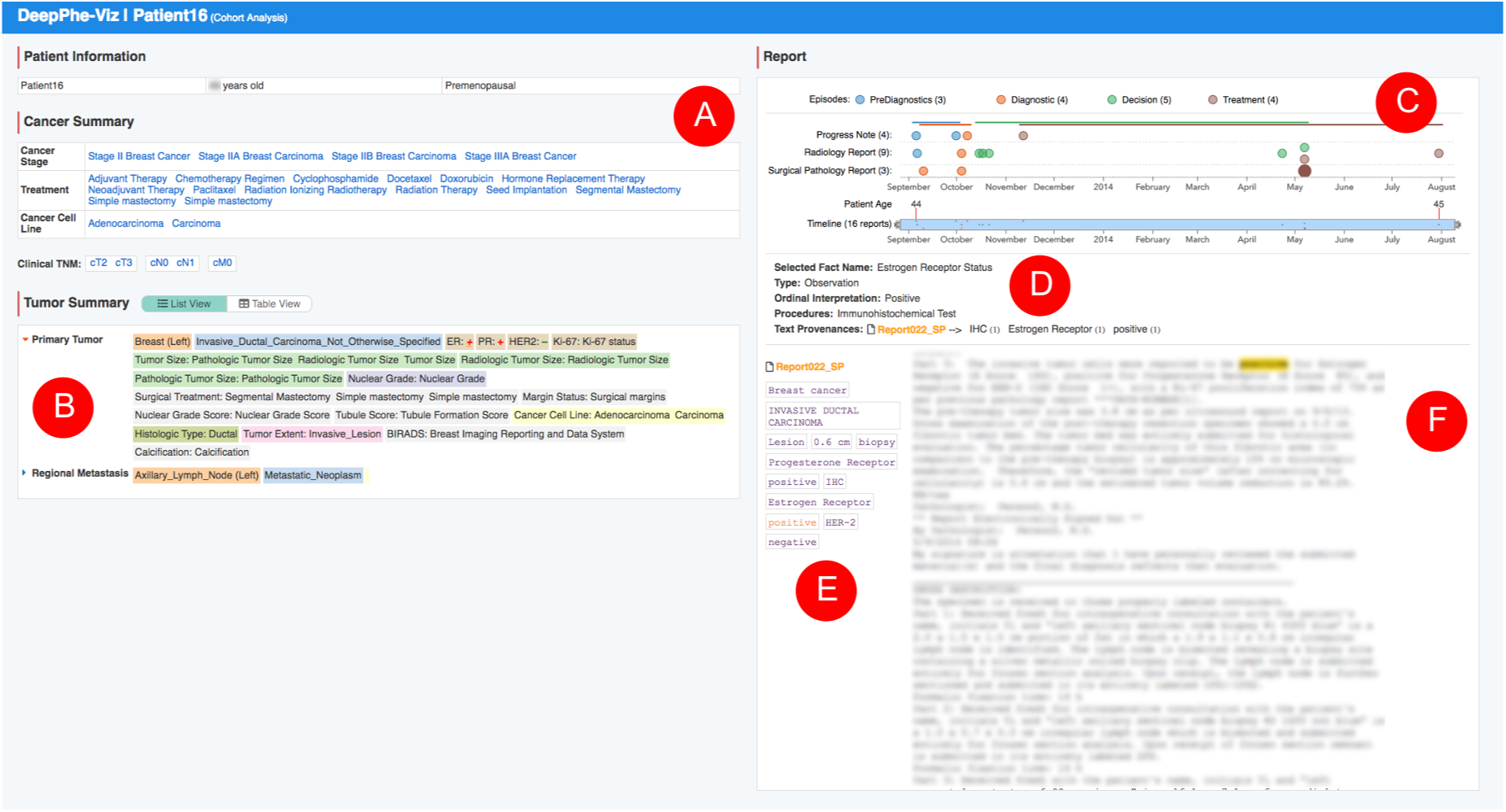
The patient visualization view: (A) Patient demographics and cancer summary. (B) Tumor summary. (C) Interactive timeline displaying documents by type and episode. (D) Inference provenance display. (E) Document-level mentions. (F) Clinical note. Text obscured to protect patient privacy.

*Tumor details* (1B) are shown as expandable lists of attributes colored to indicate classes of information (body site, diagnosis, biomarker values, etc.). As many patients have multiple tumors and information availability might vary across tumors, this approach provides a compact summary. A toggle above the tumor summary pane supports switching to tabular views when desired. Tumor and cancer details can be selected to reveal individual text spans contributing to the summary element.

The *Clinical note timeline* (1C) arranges notes on a timeline with one lane for each type of note (progress, radiology, and surgical pathology). Notes are color-coded according by episode. A double-thumb scroll-bar below the timeline allows zoom and panning across the extent, which spans from the interval between the first and last available documents. Episode labels above the timeline can be clicked to zoom the timeline to documents contained in the specified episode.

Below the timeline, the *explanation panel* (1D) bridges the gap between the inferred attributes of the cancer and tumor summaries (1A and 1B) and the text of the clinical note (1F). Selection of a summary items from the cancer or tumor summary lists leads to a display in the explanation panel describing the selected fact, along with information about its derivation from the given document.

The *mention pane* (1E) provides a summary mentions extracted from the selected document. Each mention can be clicked to highlight the appropriate scan in the *note view* (1F).

Navigation through multiple levels of abstraction is illustrated in Figure 1: The selection of tumor summary item “ER+” (1B) led to the display of the “Estrogen Receptor Status” in the explanation pane (1D), and the display of relevant mentions from Report 22 (E). Clicking on the “positive” mention leads to text confirming the ER-positive observation (1F).

The DeepPhe cohort viewer is in early stages of development. Preliminary features include selectable histograms displaying frequencies of cancer stages and biomarker status, and box plots for distribution of age at first encounter across stages, as well as distributions of stage and diagnosis information. (Figure 2).

**Figure 2:**
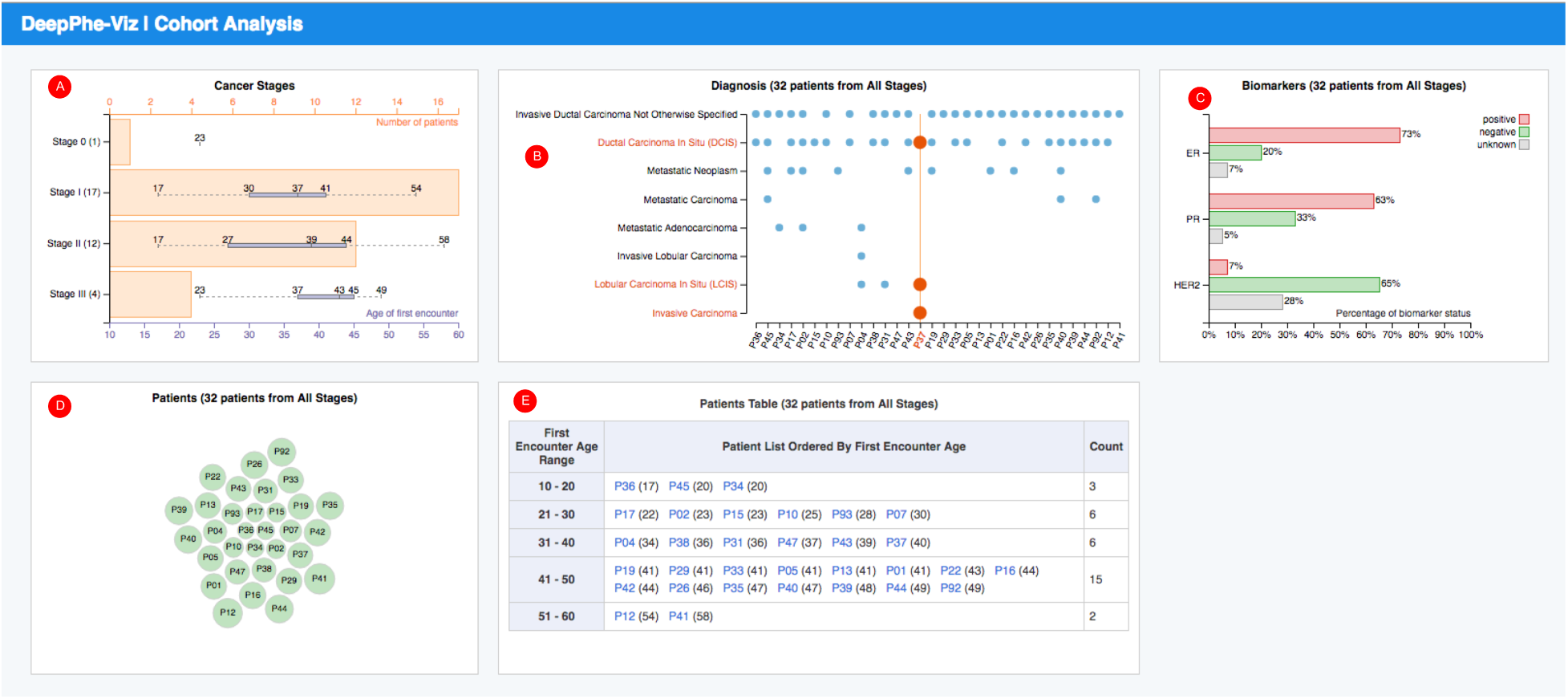
Preliminary cohort view: (A) Distributions of cancer stage and ages. (B) Diagnoses by patient, with mouse-over highlight of three diagnoses associated with the selected patient. (C) Biomarkers. (D) Bubble graph, sized by patient age at date of first note. (E) Patient list by age.

## Discussion

The substantial amounts of clinical text associate with cancer patient histories present significant challenges for retrospective research. With histories involving dozens of relevant notes (one dataset used in DeepPhe averaged 30 notes/patient, after clearly irrelevant notes were removed), manual expert review will not be sufficient for the large-scale analyses needed to drive innovation. Although advances in cross-document co-reference^15^ and other techniques currently being explored by the DeepPhe project show great promise in increasing the utility of clinical text, NLP is only a first step, providing an intermediate representation not directly consumable by end-users.

Two-key challenges must be addressed to convert concepts and relationships extracted by NLP systems into artifacts useful for clinicians and researchers. Aggregation of individual observations into higher-level clinically meaningful constructs will be necessary to easily answer key research questions such as “which patients were treated with neo-adjuvant therapy?”Once such aggregations are available, visual analytics tools will be needed to help domain experts interpret the results.

The DeepPhe visualization tools represent a first step toward these goals, providing preliminary patient-and cohort-views of cancer patient data at multiple granularities. Although limited to a subset of desired data types, the current prototype provides illustration of basic functionality needed to address key requirements (Table 2) and outstanding challenges that have been identified during the evolution of the tools.

Unlike may previous text analytics tools that focus on classification^16^ or more exploratory analysis of large text corpora^17^, the DeepPhe tools attempt to combine NLP results with an analytics interface, thus forming a complete analytics platform. DeepPhe is perhaps most similar to HARVEST^18^, which presents observation extracted from NLP in a timeline view. However, DeepPhe’s information model and inference rules provide support for higher-level abstractions not found in HARVEST.

Expanding the utility of the clinical text for both identifying cohorts and individual patients may aid in the interpretive process. Improved displays for both rendering and interpreting inference rules linking higher-order abstractions to individual text mentions may be helpful for complex inferences, particularly when cross-document inference is involved. Techniques for linking observations across documents will also prove useful for identifying recurring concepts identified through cross-document co-reference resolution. At the cohort level, visualization of text patterns, perhaps enhanced through a Word Tree^19^ or similar visualization, might help users interpret key phrases indicative of observations of interest.

DeepPhe visualization functionality will evolve alongside NLP capabilities. Although extraction and classification of individual mentions has led to promising results in many of the attributes currently shown in the prototype visualizations, much work remains to be done in the inference of higher-level aggregations, and, subsequently, the inclusion of these representations in the visualization. Two key examples involve tumors and treatments. Linking multiple tumor references across temporal extents, and including these intervals in the timeline view, will provide valuable perspective on cancer progression and response.

Enhanced temporal aggregation will also drive extensions of the DeepPhe cohort view. Incorporation of per-document episode enhancement techniques alongside orderings of treatments and time spans of specific tumors, will support temporally-aligned cohort analysis using techniques similar to those used in Outflow^8^, Frequence^10^, EventFlow^9^ and related systems^11^. Temporal^9^ and logical^20^ search facilities are also planned, with pattern search^11, 21^, a possibility for future work. Similar to previous tools focused on specific domains^22^ or care pathways and treatment plans^23–25^, we will use episode annotations and the semantics of the DeepPhe information model to focus deigns on the specific challenges of interpreting cancer data.

Inclusion of treatment information, particularly for chemotherapeutic regimens, may provide investigators with insights into treatment histories and possible impacts. Effectively displaying treatments will require inferences not only of specific start and stop times of various drugs, but ideally of identification or inference of multi-drug protocols. Extension of DeepPhe NLP tools to identify medication regimens based on the HemOnc ontology^26, 27^ is a high priority.

As DeepPhe visual analytics tools evolve to include these new data elements, appropriate handling of uncertainty and missing information will become increasingly critical. NLP temporal modeling techniques^28^, might be used in combination with structured EHR data to eliminate some ambiguity, but many details will likely remain unspecified. Cohort and patient tools will need both appropriate display of these under-specified constraints and appropriate semantics for any related queries or filters. Temporal ambiguities also underscore the importance of tools for explicitly describing search criteria explicit and for facilitating comparisons between cohorts as techniques that might reduce the risk of misinterpretation.

## Conclusion

Facilitating the extraction of understanding from complex, longitudinal patient histories is an important challenge for understanding cancer treatment and outcomes. The DeepPhe project uses a multi-faceted approach, combining both NLP, inference, information modeling, and visual analytics, to provide researchers with detailed descriptions that span the gap between key phenomena of interest and specific documentary evidence. Extension of proposed prototype designs to handle richer data, particularly involving temporal spans, will set the stage for deployment with clinical researchers and subsequent evaluation studies.

## Acknowledgments

Rebecca Jacobson, Eugene Tseytlin, Girish Chavan, and Melissa Castine made important contributions to early phases of this work. DeepPhe is supported by NCI grant 1U24CA184407.

## References

[1] Warner JL, Anick P, Hong P, Xue N. Natural Language Processing and the Oncologic History: Is There a Match? Journal of Oncology Practice. 2011;7(4):e15–e19. PMID: 22043196.

[2] AAlAbdulsalam AK, Garvin JH, Redd A, Carter ME, Sweeny C, Meystre SM. Automated Extraction and Classification of Cancer Stage Mentions from Unstructured Text Fields in a Central Cancer Registry. In: AMIA Jt Summits Transl Sci Proc; 2018. p. 16–25.

[3] Warner JL, Levy MA, Neuss MN, Warner JL, Levy MA, Neuss MN. ReCAP: Feasibility and Accuracy of Extracting Cancer Stage Information From Narrative Electronic Health Record Data. Journal of Oncology Practice. 2016;12(2):157–158.

[4] Xie F, Lee J, Munoz-Plaza C, Hahn E, Chen W. Application of text information extraction system for real-time cancer case identification in an integrated healthcare organization. Journal of Pathology Informatics. 2017;8(1):48.

[5] Kreimeyer K, Foster M, Pandey A, Arya N, Halford G, Jones SF, et al. Natural language processing systems for capturing and standardizing unstructured clinical information: A systematic review. Journal of Biomedical Informatics. 2017;73:14 – 29.

[6] Hochheiser H, Castine M, Harris D, Savova G, Jacobson RS. An information model for computable cancer phenotypes. BMC Medical Informatics and Decision Making. 2016 Sep;16(1):121.

[7] Savova GK, Tseytlin E, Finan S, Castine M, Miller T, Medvedeva O, et al. DeepPhe: A Natural Language Processing System for Extracting Cancer Phenotypes from Clinical Records. Cancer Research. 2017;77(21):e115–e118.

[8] Wongsuphasawat K, Gotz D. Exploring Flow, Factors, and Outcomes of Temporal Event Sequences with the Outflow Visualization. IEEE Transactions on Visualization and Computer Graphics. 2012 Dec;18(12):2659–2668.

[9] Monroe M, Lan R, Lee H, Plaisant C, Shneiderman B. Temporal Event Sequence Simplification. IEEE Transactions on Visualization and Computer Graphics. 2013 Dec;19(12):2227–2236.

[10] Perer A, Wang F. Frequence: Interactive Mining and Visualization of Temporal Frequent Event Sequences. In: Proceedings of the 19th International Conference on Intelligent User Interfaces. IUI ’14. New York, NY, USA: ACM; 2014. p. 153–162.

[11] Gotz D, Wang F, Perer A. A methodology for interactive mining and visual analysis of clinical event patterns using electronic health record data. Journal of Biomedical Informatics. 2014;48:148 – 159.

[12] Savova GK, Masanz JJ, Ogren PV, Zheng J, Sohn S, Kipper-Schuler KC, et al. Mayo clinical Text Analysis and Knowledge Extraction System (cTAKES): architecture, component evaluation and applications. Journal of the American Medical Informatics Association. 2010;17(5):507–513.

[13] Bostock M, Ogievetsky V, Heer J. D3 Data-Driven Documents. IEEE Transactions on Visualization and Computer Graphics. 2011 Dec;17(12):2301–2309.

[14] Gospodarowicz MK, Brierley JD, Wittekind C. TNM Classification of Malignant Tumours. John Wiley & Sons; 2017.

[15] Miller T, Dligach D, Bethard S, Lin C, Savova G. Towards generalizable entity-centric clinical coreference resolution. Journal of Biomedical Informatics. 2017;69:251 – 258.

[16] Heimerl F, Koch S, Bosch H, Ertl T. Visual Classifier Training for Text Document Retrieval. IEEE Transactions on Visualization and Computer Graphics. 2012 Dec;18(12):2839–2848.

[17] Göorg C, Liu Z, Stasko J. Reflections on the evolution of the Jigsaw visual analytics system. Information Visualization. 2014;13(4):336–345.

[18] Hirsch JS, Tanenbaum JS, Lipsky Gorman S, Liu C, Schmitz E, Hashorva D, et al. HARVEST, a longitudinal patient record summarizer. Journal of the American Medical Informatics Association. 2015;22(2):263–274.

[19] Wattenberg M, Vigas FB. The Word Tree, an Interactive Visual Concordance. IEEE Transactions on Visualization and Computer Graphics. 2008 Nov;14(6):1221–1228.

[20] Krause J, Perer A, Stavropoulos H. Supporting Iterative Cohort Construction with Visual Temporal Queries. IEEE Transactions on Visualization and Computer Graphics. 2016;22(1):91–100.

[21] Shknevsky A, Shahar Y, Moskovitch R. Consistent discovery of frequent interval-based temporal patterns in chronic patients data. Journal of Biomedical Informatics. 2017;75:83 – 95.

[22] Huang CW, Syed-Abdul S, Jian WS, Iqbal U, Nguyen PAA, Lee P, et al. A novel tool for visualizing chronic kidney disease associated polymorbidity: a 13-year cohort study in Taiwan. Journal of the American Medical Informatics Association. 2015;22(2):290–298.

[23] Aigner W, Miksch S. CareVis: Integrated visualization of computerized protocols and temporal patient data. Artificial Intelligence in Medicine. 2006;37(3):203 – 218. Knowledge-Based Data Analysis in Medicine.

[24] Bettencourt-Silva JH, Mannu GS, de la Iglesia B. Visualisation of integrated patient-centric data as pathways: Enhancing electronic medical records in clinical practice. In: Lecture Notes in Computer Science (including subseries Lecture Notes in Artificial Intelligence and Lecture Notes in Bioinformatics). vol. 9605 LNCS; 2016. p. 99–124.

[25] Gschwandtner T, Aigner W, Kaiser K, Miksch S, Seyfang A. CareCruiser: Exploring and visualizing plans, events, and effects interactively. In: IEEE Pacific Visualization Symposium 2011, PacificVis 2011 - Proceedings; 2011. p. 43–50.

[26] Malty AM, Jain SK, Yang PC, Harvey K, Warner JL. Computerized Approach to Creating a Systematic Ontology of Hematology/Oncology Regimens. JCO Clinical Cancer Informatics. 2018;p. 1–11.

[27] Warner JL, Cowan AJ, Hall AC, Yang PC. HemOnc.org: A Collaborative Online Knowledge Platform for Oncology Professionals. Journal of Oncology Practice. 2015;11(3):e336–e350. PMID: 25736385.

[28] Lin C, Dligach D, Miller TA, Bethard S, Savova GK. Multilayered temporal modeling for the clinical domain. Journal of the American Medical Informatics Association. 2016;23(2):387–395.

